# Neomorphic DNA-binding enables tumor-specific therapeutic gene expression in fusion-addicted childhood sarcoma

**DOI:** 10.1101/2022.01.05.475061

**Authors:** Tilman L. B. Hölting, Florencia Cidre-Aranaz, Dana Matzek, Bastian Popper, Severin J. Jacobi, Jing Li, Ignazio Piseddu, Bruno L. Cadilha, Stephan Ledderose, Jennifer Zwilling, Shunya Ohmura, David Anz, Annette Künkele, Frederick Klauschen, Thomas G. P. Grünewald, Maximilian M. L. Knott

**Affiliations:** Max-Eder Research Group for Pediatric Sarcoma Biology, Institute of Pathology, Faculty of Medicine, LMU Munich, Munich, Germany; Hopp Children’s Cancer Center (KiTZ), Heidelberg, Germany; Division of Translational Pediatric Sarcoma Research, German Cancer Research Center (DKFZ), German Cancer Consortium (DKTK), Heidelberg, Germany; Core Facility Animal Models, Biomedical Center, Ludwig-Maximilians-University, Planegg-Martinsried, Germany; Department of General, Visceral and Transplant Surgery, University Hospital, LMU Munich, Munich, Germany; Division of Clinical Pharmacology, Department of Medicine IV, Klinikum der Universität München, Munich, Germany; Department of Medicine II, University Hospital, Ludwig-Maximilians-Universität München, Munich, Germany; Institute of Pathology, Faculty of Medicine, LMU Munich, Munich, Germany; Charité – Universitätsmedizin Berlin, corporate member of Freie Universität Berlin and Humboldt-Universität zu Berlin, Department of Pediatric Oncology and Hematology, Augustenburger Platz 1, 13353 Berlin, Germany; Berlin Institute of Health at Charité – Universitätsmedizin Berlin, Charitéplatz 1, 10117 Berlin, Germany; German Cancer Consortium (DKTK), Berlin, Germany; German Cancer Consortium (DKTK), partner site Munich, Munich, Germany; German Cancer Research Center (DKFZ), Heidelberg, Germany; Institute of Pathology, Heidelberg University Hospital, Heidelberg, Germany

**Keywords:** Ewing sarcoma, rhabdomyosarcoma, fusion oncogene, targeted therapy, cancer gene therapy, GPR64

## Abstract

Chimeric fusion transcription factors are oncogenic hallmarks of several devastating cancer types including pediatric sarcomas, such as Ewing sarcoma (EwS) and alveolar rhabdomyosarcoma (ARMS). Despite their exquisite specificity, these driver oncogenes have been considered largely undruggable due to their lack of enzymatic activity.

Here, we show in the EwS model that – capitalizing on neomorphic DNA-binding preferences – the addiction to the respective fusion transcription factor EWSR1-FLI1 can be leveraged to express therapeutic genes.

We genetically engineered a *de novo* enhancer-based, synthetic and highly potent expression cassette that can elicit EWSR1-FLI1-dependent expression of a therapeutic payload as evidenced by episomal and CRISPR-edited genomic reporter assays. Combining *in silico* screens and immunohistochemistry, we identified GPR64 as a highly specific cell surface antigen for targeted transduction strategies in EwS. Functional experiments demonstrated that anti-GPR64-pseudotyped lentivirus harboring our expression cassette can specifically transduce EwS cells to promote the expression of viral thymidine kinase sensitizing EwS for treatment to the otherwise relatively non-toxic (Val)ganciclovir and leading to strong anti-tumorigenic, but no adverse effects *in vivo*. Further, we prove that similar vector designs can be applied in PAX3-FOXO1-driven ARMS, and to express immunomodulatory cytokines, such as IL-15 and XCL1, in tumor types typically considered to be immunologically ‘cold’.

Collectively, these results generated in pediatric sarcomas indicate that exploiting, rather than suppressing, the neomorphic functions of chimeric transcription factors may open inroads to innovative and personalized therapies, and that our highly versatile approach may be translatable to other cancers addicted to oncogenic transcription factors with unique DNA-binding properties.

## Background

Unlike most malignancies in adults, childhood sarcomas are commonly characterized by a striking paucity of somatic mutations^1^. However, these entities often harbor tumor-defining fusion oncogenes, such as *EWSR1-FLI1* (EF1) in Ewing sarcoma (EwS) and *PAX3-FOXO1* (P3F1) in alveolar rhabdomyosarcoma (ARMS) acting as potent drivers of malignancy^2,3^. Both chimeric oncogenes exert their function as aberrant transcription factors equipped with neomorphic features allowing them to bind unique DNA motifs that differ from the binding sites of their parental constituents^4,5^. For example, EF1 binds to otherwise non-functional GGAA-microsatellites (msats), which are thereby converted into potent *de novo* enhancers^6^. Even though the interaction between EF1 and GGAA-msats is incompletely understood, accumulating evidence suggests that EF1 preferentially binds to GGAA-msats with a specific structure (min. 4 GGAA-repeats; optimal binding at 15–25 GGAA-repeats)^4,7^. Similarly, P3F1 binds to a highly specific motif (ATTWGTCACGGT), which induces disease-defining, myogenic super enhancers^5,8^. In both cancer types, these aberrant DNA binding preferences of the respective chimeric oncoproteins massively deregulate the cellular transcriptome, which promotes their malignant phenotype and oncogene-addiction^5,9^.

Based on the specificity of their interaction with fusion transcription factors and the oncogene-dependency exhibited by the tumors expressing these oncoproteins, we hypothesized that these aberrantly bound neo-enhancers would represent ideal candidates to drive a tumor-specific expression of therapeutic genes.

## Methods

### Provenience of cell lines, and cell culture conditions

Human cell lines 293T, HeLa, Hep-G2, MHH-ES1, PA-TU-8988T, RD-ES, RH30, SK-N-MC, and U2-OS were obtained from the German Collection of Microorganisms and Cell Cultures (DSMZ) (Braunschweig, Germany). The human A-673 and MRC-5 cell lines were purchased from the American Type Culture Collection (ATCC). TC-71, TC-106 was obtained from the Children’s Oncology Group (COG). RH4 and RD were kindly gifted by R. Kappler (Munich, Germany). A-673/TR/shEF1 and A-673/TR/shctrl were kindly gifted by J. Alonso (Madrid, Spain)^10^. Primary human umbilical vein endothelial cells (HUVEC) were kindly provided by S. Massberg (Munich, Germany). All cell lines were cultivated in a humidified atmosphere with 5% CO_2_ at temperature of 37°C. Except for 293T and HUVEC, all cell lines were grown in RPMI-1640 medium containing stable L-glutamine and sodium bicarbonate (Sigma-Aldrich) supplemented with 10% tetracycline-free fetal bovine serum (FCS) (Sigma-Aldrich). Unless specified otherwise, 293T were grown in DMEM supplemented with 10% FCS (Sigma-Aldrich). HUVEC were cultured in Endothelial Cell Growth Medium (Cell Applications, Inc). Cell lines were routinely tested for mycoplasma contamination by nested PCR, and cell line identity was routinely confirmed by STR profiling.

### Analysis of published chromatin-immunoprecipitation and sequencing (ChIP-seq) and RNA-sequencing (RNA-seq) data

Raw data of published ChIP-seq experiments of EF1 in Ewing sarcoma cell lines A-673 and SK-N-MC were retrieved from the European Nucleotide Archive (Samples used: SRR1593960, SRR1593966, SRR1593985, SRR1593991)^6^. Data analysis was performed following the ENCODE Transcription Factor and Histone ChIP-seq processing pipeline^11^: First, quality of raw data was assessed with FastQC^12^. Reads were aligned to the human reference genome (*hg38)* using *bowtie2*^13^. Alignments that were unmapped, not primarily aligned, failing platform checks, duplicates, or aligned with c <30 were removed using s*amtools view* and *picard MarkDuplicates*^14,15^. Peaks were called on by *SPP* from the bioconda package *phantompeakqualtools* using npeak = 30000^11^. To determine the peaks reproducible for both A-673 and SK-N-MC, the Irreproducibility Discovery Rate (IDR) framework was applied with a threshold of 5% using the bioconda package *idr*^16^. Genomic locations (*hg38*) of GGAA-msats, defined as 4 GGAA repeats on either strand, were obtained using the *locate* function of *seqkit*^4,17^. GGAA-msats overlapping or within 100bp from each other, were merged with the *merge* function of the bedtools suite^18^. By intersecting the EF1 peaks with the GGAA-msat locations, EF1-bound msats were defined, and their genomic annotation and closest RefSeq TSS were obtained using HOMER^18,19^. For analysis of previously published RNA-seq experiments from the same study were retrieved from the European Nucleotide Archive (samples used: SRR1594020, SRR1594021, SRR1594024, SRR1594025). *salmon* was used to quantify transcript abundance^20^. Differentially expressed genes after 48h of shRNA-mediated knockdown of EF1 compared to control cell lines without knockdown were determined by use of the R package *DeSeq2*^21^. For a more conservative estimation for genes with low read counts, log2FCs were shrunk using *apeglm* within the *lcfShrink* wrapper function^22^.

### Cloning and plasmid preparation

All cloning was designed on benchling.com and performed using standard restriction-ligation approaches. Plasmid integrity was verified using Sanger-sequencing and agarose gel electrophoresis. All primers and sequences of synthetic linear DNA molecules used for cloning can be found in **Additional Table 4**. To create firefly luciferase reporter plasmids, we inserted a PacI restriction site into the multiple cloning site (MCS) of pGL4.17 [*luc2*/Neo] (Promega) upstream of the EcoRV restriction site using annealed oligonucleotides (Eurofins Genomics) following a KpnI-EcoRV restriction digest. We then inserted double stranded synthetic DNA fragments composed of 17, 21, or 25 GGAA-repeats upstream of the recently described minimal activity promoter YB-TATA (Genscript) into the newly designed MCS by PacI-EcoRV restriction digest^23^.

Therapeutic plasmids were sequentially created using the EF.CMV.RFP backbone (Addgene #17619) by replacing the EF1 promoter by the synthesized 25-GGAA-YB-TATA fragment (Genscript) followed by a firefly *luciferase* (cloned from pcDNA3-luciferase (Addgene #18964)) or *HSV-TKSR39* (cloned from pET23d:HSVTK-SR39 (kindly provided by Margaret Black, Washington State University)) gene or both P2A-fused genes, and a WPRE (cloned from pLenti CMV GFP Puro (658-5) (Addgene #17448)) all flanked by epigenetic insulator sequences by repetitive restriction digests (KflI-EcoRI, PacI-EcoR1, SpeI-EcoRI, SalI-EcoRI, Bsu36I-AsiSI). RFP was replaced by the puromycin-resistance gene (cloned from pLenti CMV GFP Puro (658-5)), respectively, using a BamHI-NsiI restriction digest after additional insertion of a NsiI digestion site by PCR. Lastly, the full construct was transferred to the p156RRL-sinPPT backbone derived from pLenti CMV GFP Puro (658-5) after ClaI-KpnI restriction. To this end, we inserted a new MCS harboring an additional AgeI restriction site using annealed oligonucleotides (Eurofins Genomics) and inserted the construct by AgeI-NsiI restriction digest resulting in *pLenti_25_LT_Puro*. For creation of all other plasmids, insulator sequences were removed by PCR. For IL-15 and XCL1 expressing plasmids, the *IL-15*-P2A-*XCL1* gene was synthesized by Genscript and inserted by SpeI-SalI digest. To allow better secretion, a mouse IgV signal peptide was inserted right after the start codon of IL-15^24^. To reduce the toxicity of XCL1 on *E. coli*, a synthetic intron (113 nt) was included^25^. For genomic reporter assays, the microsatellite-YB-TATA fragment was excised by PacI-SpeI double restriction digest and replaced by a synthetic, unique gRNA binding sequence from the *luciferase* gene fused to YB-TATA. Subsequently, the *HSV-TKSR39*-P2A-*luciferase* transgene was replaced by eGFP cloned from pLenti CMV GFP Puro (658-5) by SpeI-SalI restriction digest. gRNA binding prediction was performed using the CRISPR tool of benchling.com. For ARMS, the GGAA-microsatellite was replaced by genomic or synthetic alk-super-enhancer (SE) sequences fused to the YB-TATA promoter by double restriction digest (SbfI-SpeI)^5^. Full plasmid maps can be retrieved from the authors upon request. All ligation reactions were performed at room temperature (RT) for 30 min using T4 ligase (NEB). Bacterial transformation was performed following standard protocols using Stellar competent cells (Takara). Mini- and MidiPreps (Macherey-Nagel) were performed according to the manufacturer’s instructions.

### Extraction of total DNA and RNA, reverse transcription, and quantitative Real-Time PCR (qRT-PCR)

Total DNA extraction was performed using NucleoSpin® Tissue mini kit (Macherey-Nagel, Germany). RNA was extracted using NucleoSpin® RNA mini kit (Macherey-Nagel, Germany) including a 15 min DNAse-treatment, and reversely transcribed using the High-Capacity cDNA Reverse Transcription Kit (Applied Biosystems). qRT-PCRs were performed in a final volume of 15 µl using SYBR™ Select Master Mix (Applied Biosystems). All primer sequences used for qRT-PCR are listed in **Additional Table 4**. Cycling conditions were as follows: 95°C for 10 min (initial denaturation), then 50 cycles at 95°C for 10 sec (denaturation) and 60°C for 1 minute (annealing, elongation and detection).

### Lentivirus production and concentration

Lentivirus production was performed by polyethylenimine (PEI)-mediated transfection of adherent 293T cells as previously described^26^. In short, 24 h before transfection, 5.3×10^5^ cells were seeded per well (6-well) in 2 ml fully supplemented DMEM containing 10% FCS. On day of transfection 1020 ng lentiviral transfer plasmid, 680 ng of pCD/NL-BH*DDD (Addgene plasmid # 17531) and either 340 ng of pCEF-VSV-G (Addgene plasmid # 41792) or 680ng of 2.2 (Addgene plasmid # 34885) were mixed in 100 µl final volume of Opti-MEM (Gibco). 15.12 µl of PEI Max Transfection Grade Linear Polyethylenimine Hydrochloride MW 40.000 (Polysciences) (1 mg/ml) were diluted in a final volume of 100 µl Opti-MEM in a separate tube. After 5 min of individual incubation, the PEI mix was added to the plasmid mix and mixed by pipetting. The resulting PEI-plasmid mix was incubated for 5 min. In the meantime, the medium of the previously seeded 293T cells was replaced by 2 ml of, either DMEM for transfections using VSV-G virus, or UltraCULTURE medium (Lonza) for transfections with 2.2. After incubation, the PEI-plasmid mix was added to the medium dropwise. The medium was replaced by fresh DMEM or UltraCULTURE 16 h after transfection. After an additional 48 h, the supernatant containing the viral particles was collected. To remove cellular debris, the harvested medium was centrifuged for 5 min at 1000 g and the resulting supernatant filtered through a syringe filter (0.45 µm pore size, CA membrane). When high viral titers were needed (i.e. for *in viv*o experiments), virus production was upscaled on 150 mm dishes and the viral supernatant was concentrated using polyethylene glycol-based precipitation. To this end, 3 parts of supernatant were mixed with 1 part of custom-made lentivirus concentrator solution (40% w/w PEG8000, 1.337 M NaCl, 2.7 mM KCl, 8 mM Na_2_HPO_4_ und 2 mM KH_2_PO_4_) and incubated at 4°C overnight on a roller mixer. After 24 h, the mixture was centrifuged at 2000 g for 1 h, the supernatant was discarded and the resulting pellets were resuspended in PBS. Viral titers were calculated based on flow cytometry (FACSCanto™ II, BD) results or qPCR as described previously^26^.

### *In vitro* lentiviral transduction

Cell lines were transduced using a standard MOI of 2 (U2-OS reference) by adding equal amounts of virus-containing, filtered supernatants as previously described^27^. Where indicated, cells were selected using puromycin (Invivogen) at the minimum concentration necessary for complete cell death determined for each cell line individually.

### Clustered regularly interspaced short palindromic repeats (CRISPR) knock-in

For genomic reporter assays, A-673 cells were lentivirally transduced with a *GFP* reporter gene downstream of the minimal promoter YB-TATA^23^ and a unique guide-RNA binding site. After puromycin-selection, the transduced cells were single cell cloned and clones harboring a single copy of the reporter construct were identified using genomic qPCR as previously described^26^. Two clones were selected and 4×10^4^ cells in 100 µl complete medium were CRISPRed using reverse lipofection technique (Lipofectamine CRISPRMAX, LifeTechnologies) of 30 nM ALT-R CAS9 nucleoproteins (IDT) and 25 nM 25 GGAA-repeat HDR templates (50 bp homology arms each, IDT) in a 96-well plate. After 48 h, the medium was changed and the cells were expanded. GFP-positive cells were single-cell-sorted into a 96-well plate using a FACSAria™ II (BD). After expansion of the individual GFP-positive clones, cells were lysed and DNA was isolated as described above. The correct insertion of 25 GGAA-repeats upstream of the YB-TATA promoter was confirmed by Sanger-sequencing of the PCR-amplified genomic region. Cells were imaged using a Zeiss Axiovert 25 microscope and the Zeiss AxioVision (Release 4.9.1 SP2) software. Images were overlaid and contrast-enhanced using Adobe Photoshop (Adobe). Fluorescence intensity was measured by flow cytometry (FACSCanto™ II, BD).

### Dual-luciferase reporter assays

5×10^4^ cells were seeded per well (24-well) 24 h prior to transfection. Cells were co-transfected with the plasmid pGL4.17 containing a YB-TATA-based minimal activity promoter and enhancer sequences (GGAA-msats of various lengths, genomic or synthetic alk-SE sequences) and pRL *Renilla* Luciferase Control Vector (Promega) (plasmid mass ratio of pGL4.17:pRL = 100:1). Where indicated, reporter plasmids were co-transfected with a plasmid containing either *EWSR1-FLI1* cDNA (pCDH-CMV-E/F1-puro) or a defective mutant (or pCDH-CMV-E/F1_R2L2-puro^28^). Furthermore, in experiments with A673/TR/shEF1 and A673/TR/shCtrl medium was supplemented with doxycycline 1 µg/ml (Sigma-Aldrich) to achieve shRNA-mediated knockdown of EF1. Transfections were performed using PEI MAX (Polysciences) for all cell lines apart from Jurkat and RD for which Lipofectamine LTX was used. 12 h after transfection, the medium was replaced with fresh RPMI containing 10% FCS. After 36 h, cells were lysed and luminescence measured using a dual-luciferase assay kit (Beetle-Juice Luciferase assay firefly and *Renilla*-Juice Luciferase Assay, PJK GmbH). Firefly luciferase induced luminescence was normalized on *Renilla* luciferase luminescence and the resulting ratios were normalized to that of the empty control plasmid condition.

### Western blot

Western blots were performed as previously described^29^. For preparation of protein lysates, 3×10^5^ *pLenti_25_LT_Puro-*transduced and selected cells were seeded per well (6-well). After 48 h, medium was removed, cells were washed with 1 ml of PBS and lysed by adding 100 µl of lysis buffer containing 150 mM NaCl, 0.1% Triton X-100, 50 mM Tris-HCl at pH 8.0 supplemented with cOmplete, Mini, EDTA-free Protease Inhibitor Cocktail (Roche). Detection of specific bands for firefly luciferase or GAPDH was performed using a HRP-conjugated monoclonal Anti-Luciferase antibody (sc-74548 HRP, 1:2,000, Santa-Cruz) and a HRP-conjugated, monoclonal murine Anti-GAPDH antibody (HRP-60004, 1:50,000, Proteintech).

### Cell viability assays

5×10^3^ (1×10^4^ for TC-106) *pLenti_25_LT_Puro-*transduced and selected cells were seeded in 90 µL medium per well (96-well). After 24 h, ganciclovir (GCV) was added for final concentrations ranging from 0.01 µM to 50 µM with 0.05% dimethyl sulfoxide (DMSO) in all conditions. 72 h after the addition of GCV, cell viability was assessed using a resazurin-based readout system^30^. Relative fluorescence units of treated wells were background corrected and normalized to vehicle controls.

### Apoptosis assays

5×10^4^ of *pLenti_25_LT_Puro-*transduced and selected cells were seeded per well (24-well). 24 h later, GCV was added for final concentrations of 0.4 µM. After 72 h, apoptosis was analyzed by Annexin V/PI staining (APC Annexin V Apoptosis Detection Kit with PI, Biolegend) and flow cytometry using a FACSCanto™ II (BD) cytometer. An example of the gating strategy is found in **Additional Figure 6a**.

### IL-15 and XCL1 ELISAs

3×10^5^ *pLenti_25_IX_Puro-*transduced or wildtype cells were seeded in 0.5 ml per well (12-well) and incubated for 72 h in RPMI 1640 containing 10% FCS. Supernatants were harvested and stored at -20°C until further use. IL-15 and XCL1 levels were quantified using human IL-15 Duoset ELISA (RnD) and human XCL1 Duoset ELISA (RnD) according to the manufacturer’s protocol. Cytokine levels were calculated by constructing a 4-parameter logistic regression model based on standard measurements for each plate using the R package *drc*^31^.

### *In vitro* T cell migration assays

Migration assays were performed using 96-Transwell plates with 3 µm pore size (Corning). 225 µl conditioned medium was transferred into the lower chamber, before 1×10^6^ freshly isolated Peripheral blood mononuclear cells (PBMC) of healthy donors were loaded onto the membrane in 70 µl complete medium. After 4 h, the transwell insert was removed and the cells in the lower chamber were collected, stained for CD3 (Biolegend), analyzed by flow cytometry (FACSCanto™ II, BD) and quantified using Precision Count Beads (Biolegend). An example of the gating strategy can be found in **Additional Figure 6b**.

### Microarray analysis

To identify genes encoding membrane proteins that could serve as potential binding points for targeted transduction of EwS cells, we took advantage of a gene expression previously described data set of publicly available microarray data (Affymetrix HG-U133Plus2.0) consisting of 928 normal human tissue samples and 50 EwS samples^32^. Robust Multiarray Average (RMA) normalization and calculation of expression measures was performed using the function *just.RMA* of the bioconductor package *affy*^33^. Genes that were statistically significantly overexpressed in EwS compared to every other tissue with a fold change of at least 2 were identified using the R package *limma* using a false discovery rate (FDR) cutoff of 5%^34^. Multiple testing was accounted for using the Benjamini-Hochberg procedure. Accession codes of samples used in the analysis can be found in **Additional Table 5**.

### Analysis of protein expression by indirect flow cytometry

2×10^5^ cells were seeded per well (12-well) 24 h prior to analysis. Cells were harvested using trypsin and washed in PBS twice. Subsequently, cells were then stained with the primary antibody for the indicated antigens (0.25 µg per 1×10^6^ cells) for 30 min at RT. Cells were washed three times with PBS, before the secondary antibody (0.375 µg per 1×10^6^ cells) was applied for 30 min at RT. After three additional washing cycles, stained cells were co-stained with propidium iodide (PI) solution and analyzed on a FACSCanto™ II (BD) cytometer. An example of the gating strategy is found in **Additional Figure 6c**. The following antibodies were used: CD99 (3B2/TA8, Biolegend), FAT4 (NBP1-78381, Novus Biologicals), GPR64 (purified using Mouse TCS Antibody Purification Kit (ab128749, abcam) from OAM6#93 (PTA-5704, ATCC), GD2 (TAB-731, Creative Biolabs), Mouse IgG2b Isotype Control (#02-6300, Invitrogen), Rabbit IgG Isotype Control (#02-6102), Goat anti-Rabbit IgG (H+L) Cross-Adsorbed Secondary Antibody, APC (A-10931, Invitrogen), Goat anti-Mouse IgG (H+L) Cross-Adsorbed Secondary Antibody, APC (A-865, Invitrogen).

### Tissue microarrays and evaluation for immunoreactivity

Formalin-fixed samples tissue microarrays of EwS-samples and normal tissues were stained for GPR64 after antigen retrieval using Target Retrieval Solution (Fa.Agilent Technologies, S1699) with anti-GPR64 (purified using Mouse TCS Antibody Purification Kit (ab128749, abcam) from OAM6#93 (PTA-5704, ATCC) with a concentration of 40 µg/ml for 1 h at RT. For signal detection the MACH 3 Mouse HRP Polymer Detection system was employed according to manufacturer’s protocol using DAB+ (Fa.Agilent Technologies, K3468). Slides were counter-stained with Hematoxylin Gill’s Formula (Fa.Vector, H-3401). Signal intensities were evaluated by two blinded resident pathologists using a semi-quantitative score in analogy to the previously described Immune Reactive Score (IRS)^32^.

### Evaluation of targeted transduction *in vitro*

5×10^4^ cells were plated in 400 µl per well (24-well). 24 h later, 100 µl of unconcentrated lentiviral supernatant were mixed with 0.5 µg antibody and added to each well (final concentration of antibody: 1 µg/µl). After 24 h, supernatants were removed and cells were incubated for an additional 24 h. Cells were then harvested and fluorescence was analyzed by flow cytometry. An example of the gating strategy is found in **Additional Figure 6d**.

### *In vitro* therapy assays

5×10^3^ cells were seeded in 90 µl of medium per well (96-well). After 24 h, equal amounts of concentrated lentivirus (approx. 1000 transducing units, TU) were added to each well. After additional 24 h, GCV was added for a final concentration of 20 µM. Cell viability was assessed by a resazurin-based assay 72 h after the addition of GCV.

### Luciferase-based evaluation of promoter activity and specificity *in vivo*

To assess promoter-dependent gene expression in various non-EwS tissues 2×10^7^ TU (transducing units) of VSV-G-pseudotyped virus produced with *pLenti_25_LT* or *pLenti_CMV_LG* were injected in 200 µl PBS intraperitoneally into NSG mice (NOD.Cg-Prkdc^SCID^Il2rg^tm1Wjl^/SzJ, Charles River Laboratories). 7 days later, luminescence was measured on an IVIS-100 (Perkin-Elmer) imaging system after intraperitoneal injection of 3 mg D-luciferin.

### Mouse xenograft experiments

For subcutaneous xenograft experiments, six- to eight-week old NSG mice were subcutaneously injected into the flank with 2×10^6^ wildtype or pre-transduced A-673 or RD-ES EwS cells in Cultrex Basement Membrane Extract (R&D Systems) to enhance tumor formation. Tumor growth was measured three times a week using a caliper. Tumor volumes were calculated using the following formula: V = L × W^2^ / 2, where V is tumor volume, L is largest diameter and W smallest diameter. For intraperitoneal engraftment, 2×10^6^ luciferase-expressing A-673 cells were intraperitoneally injected. Animal experiments were approved by the government of Upper Bavaria and conducted in accordance with ARRIVE guidelines, recommendations of the European Community (86/609/EEC), and UKCCCR (guidelines for the welfare and use of animals in cancer research).

### *In vivo* tumor transduction

For the evaluation of antibody-directed transduction of subcutaneous xenografts (**Fig. 3f**), 0.5×10^6^ TU of 2.2 or VSV-G pseudotyped virus was injected intratumorally in 100 µl PBS containing 15 µg/ml antibody where indicated. For therapeutic transduction of subcutaneous xenografts (**Fig. 4a**), 1×10^7^ TU of GPR64-directed 2.2 pseudotyped lentivirus was injected intratumorally in 100 µl PBS containing 15 µg/ml anti-GPR64 antibody twice per week once the tumor had reached a mean diameter of 5 mm. For therapeutic transduction of intraperitoneal tumor masses, 2×10^7^ TU of GPR64-directed 2.2 pseudotyped lentivirus was injected intraperitoneally in 200 µl PBS containing 15 µg/ml anti-GPR64 antibody on three consecutive days (day 3 to 5), three days after tumor inoculation. A second round of three consecutive viral injections was performed on day 13 to 15. Bioluminescence imaging after intraperitoneal injection of 3 mg D-luciferin was performed on days 6, 12 and 19.

### Oral Valganciclovir (VGCV) administration

For the treatment of *pLenti_25_LT* or *pLenti_25_TK* (pre-)transduced xenografts, VGCV (0.5 mg/ml) was administered orally *ad libidum* by addition to the drinking water. To mitigate any adverse taste, 5% sucrose (Carl Roth) was added as well. 5% sucrose containing drinking water served as a control where indicated.

### Statistical analysis

Data was analyzed using R (R version 4.1.2, R Foundation for Statistical Computing, Vienna, Austria). Where not otherwise specified, the statistical significance of differences between two experimental groups were tested using the two-tailed Wilcoxon Rank Sum / Mann-Whitney test with the Holm–Bonferroni method to account for multiple comparisons; * :p <= 0.05, **: p <= 0.01, ***: p <= 0.001, ****: p <= 0.0001.

## Results

### Synthetic msat-promoter designs are functional and allow EF1-dependent gene expression

Since the neomorphic DNA-binding preferences of EF1 are very well characterized, we first turned to EwS as a model disease^35–39^. Although reanalysis of publicly available ChIP-seq data generated from two EwS cell lines (A-673, SK-N-MC) demonstrated that most (97.7%) EF1-bound GGAA-msats (defined as at least 4 consecutive GGAA-repeats) were located in intergenic and intronic regions, we identified 4 EF1-bound GGAA-msats located in direct proximity (defined as -1,000 bp to +100 bp distance) of the transcriptional start site (TSS) of genes annotated in RefSeq (0.4%) (**Fig. 1a, Additional Table 1**)^4,6^. Among those, the shortest interval between the identified EF1-bound GGAA-msat and the respective TSS (interval = 49 bp) corresponded to the lncRNA *FEZF1-AS1*, which reanalysis of published RNAseq data showed to be significantly downregulated after shRNA-mediated knockdown of EF1 (**Additional Fig. 1a and 1b**). Hence, we assumed that even minimal distance between these EF1-bound neo-enhancers and TSS doesn’t abrogate the transactivating function of EF1.

**Fig. 1:**
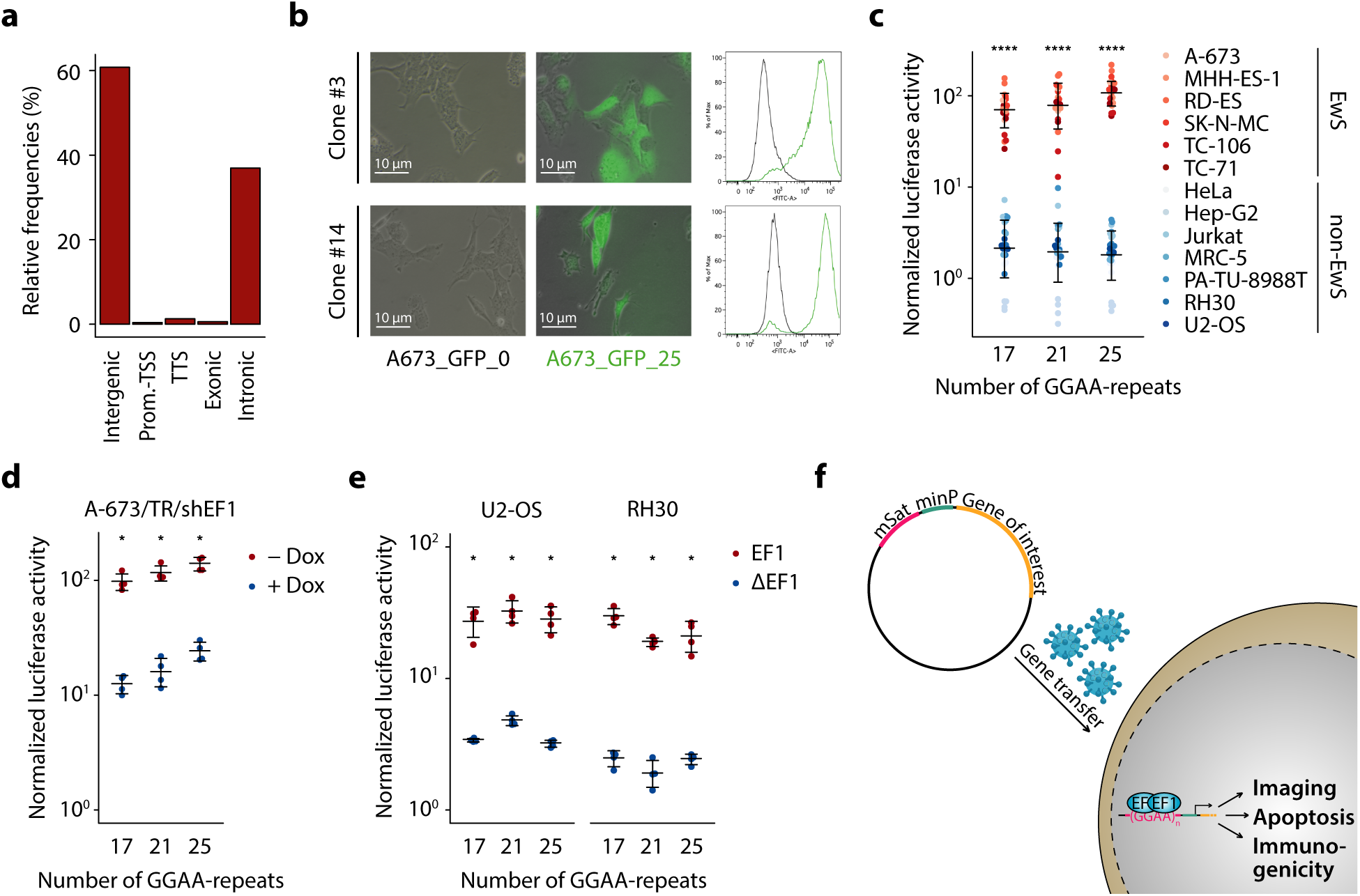
GGAA-msats allow EwS-specific and EF1-dependent gene expression. **a** Genomic annotation of location of EF1-bound GGAA-msats (min. 4 GGAA-repeats). **b** Fluorescent microscopy image (left) and flow cytometry histograms (right) of A-673 stably transduced with GFP under the control of a minimal promoter and with or without CRISPR/Cas9-mediated knock-in of 25 GGAA-repeats (A673_GFP_25 / A673_GFP_0) in two independent single cell clones. **c** Luciferase reporter assays of indicated EwS and non-EwS cell lines after co-transfection with a reporter plasmid containing the indicated number of GGAA-repeats upstream of the minimal promoter YB-TATA and a constitutively expressed *Renilla*-encoding plasmid. Dots indicate firefly to *Renilla* luminescence ratios normalized to a reporter plasmid without GGAA-repeats for 4 biologically independent experiments. Horizontal bars indicate mean and whiskers standard deviation per group. **d** Luciferase reporter assays of A-673/TR/shEF1 co-transfected with the same plasmids as in Fig. 1c treated with / without Dox. Dots indicate firefly to *Renilla* luminescence ratios normalized to a reporter plasmid without GGAA-repeats for 4 biologically independent experiments. Horizontal bars indicate mean and whiskers standard deviation per group. **e** Luciferase reporter assays of non-EwS cell lines RH30 and HeLa co-transfected with the same plasmids as in Fig. 1c and a plasmid expressing EF1 or a defective mutant of EF1 (ΔEF1). Dots indicate firefly to *Renilla* luminescence ratios normalized to a reporter plasmid without GGAA-repeats for 4 biologically independent experiments. Horizontal bars indicate mean and whiskers standard deviation per group. **f** Schematic representation of EF1-dependent expression cassette and potential use. P values were determined with two-tailed Mann-Whitney test, *: p <= 0.05, ****: p <= 0.0001.

To test this hypothesis, we generated a EwS reporter cell line, in which we inserted a GGAA-msat directly upstream of a synthetic minimal promoter (YB-TATA) by CRISPR-mediated homology-directed repair (HDR)^23^. Indeed, these clones showed a strong and persistent overexpression of the reporter gene *GFP*, which was not observed in clones lacking the GGAA-msat (**Fig. 1b**). Thus, the transactivating functionality of EF1 appears to be retained when the GGAA-msat-enhancer is located closely to the respective promoter.

Prior reports have demonstrated that the affinity of EF1 to GGAA-msats, and thereby their enhancer activity, correlates positively with the number of consecutive GGAA-repeats^4,7^. We therefore tested three different expression cassettes consisting of 17, 21, or 25 GGAA-repeats cloned directly upstream of YB-TATA in a dual luciferase reporter assay in 6 EwS cell lines (including TC-106, a cell line harboring the less common *EWSR1-ERG* fusion oncogene, which is structurally and functionally similar to EF1) and 7 control cell lines, comprising 7 different non-EwS cancer entities or tissue types^1,40^. Excitingly, we observed a very strong and length-dependent induction by the evaluated GGAA-msats in all tested EwS cell lines whereas there was only minimal induction of reporter activity in the transfected control cell lines (**Fig. 1c**). To further assess the EF1-dependency of firefly luciferase expression, we repeated these reporter assays in a EwS cell line harboring a doxycycline-inducible shRNA targeting EF1 (A-673/TR/shEF1) or a control shRNA (A-673/TR/shCtrl). Strikingly, conditional knockdown of EF1 dramatically reduced the reporter signal (**Fig. 1d, Additional Fig. 1c-d**). Conversely, ectopic expression of *EF1* in non-EwS osteosarcoma and rhabdomyosarcoma cells (U2-OS, RH30) induced the reporter signal while expression of a mutant *EF1* lacking its DNA-binding capacity (EF1_mut R2L2 (ΔEF1)^28^) showed no relevant induction (**Fig. 1e**). These results strongly suggest that functional EF1 (or EWSR1-ERG) is necessary and sufficient for induction of this expression cassette in an episomal setting.

Based on these data, we reasoned that combining a 25 GGAA-repeat element and a minimal promoter could serve as a backbone to mediate the EF1-dependent expression of any therapeutic gene for targeted therapy of EwS (**Fig. 1f**).

### Synthetic msat-promoter designs are highly specific and enable therapeutic gene expression

To test this hypothesis *in vitro*, we generated a lentiviral transfer plasmid (*pLenti_25_LT_Puro*), containing these regulatory elements followed by the gene encoding a modified *Herpes simplex virus* thymidine kinase *(HSV-TK SR39*) coupled with a firefly luciferase by a P2A linker peptide^41^. We chose *HSV-TK* as a first candidate gene due to its well characterized phenotype and clinical use as suicide-gene in CAR T cell-based therapies^42^. Next, EwS and non-EwS control cell lines were transduced using this vector or an identical control vector lacking the 25 GGAA-repeats (*pLenti_0_LT_Puro)*. Successfully transduced cell lines were selected by puromycin and subjected to reverse transcription qPCR analysis for induction of *HSV-TK* transcription. EwS cell lines showed a significant induction of *HSV-TK* using the *pLenti_25_LT_Puro* vector compared to the control vector without the GGAA-repeats (*pLenti_0_LT_Puro*), whereas in the non-EwS control cell lines the expression levels were similar for both vectors (**Additional Fig. 2a**). In agreement with these findings at the mRNA level, immunoblotting confirmed that transgenes encoded by *pLenti_25_LT_Puro* were only detectable in EwS cells but not in non-EwS cells at the protein level ( **Fig. 2a**). As single cell lines do not reflect the complexity of tissues or organisms, we sought to evaluate the specificity of our expression cassette *in vivo*. To this end, we generated a transfer plasmid (*pLenti_25_LT*) similar to *pLenti_25_LT_Puro* but lacking the puromycin resistance cassette and intraperitoneally injected 1×10^7^ TU of VSV-G pseudotyped lentiviral particles carrying either *pLenti_25_LT* or a CMV-driven *luciferase* (*pLenti_CMV_LG*). Excitingly, no luciferase signal was detected in the *pLenti_25_LT* group, whereas strong luciferase signal was obtained in the thoracoabdominal region of *pLenti_CMV_LG-*transduced animals (**Fig. 2b**). To exclude differences in transduction efficiency, we harvested the organs and found comparable copy numbers of both vectors by genomic qPCR (**Additional Fig. 2b**).

**Fig. 2:**
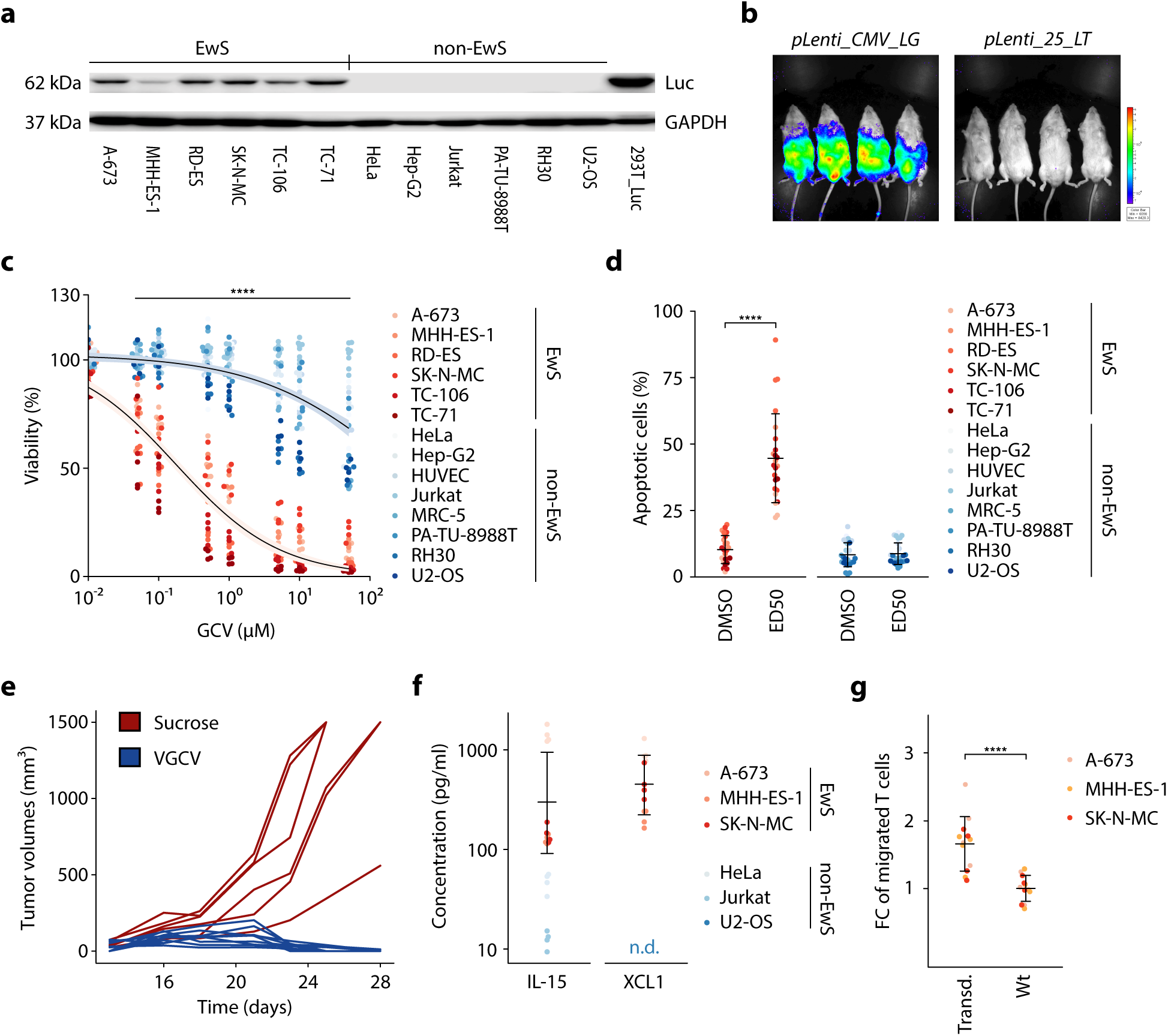
Synthetic msat-promoters enable specific and effective therapeutic gene expression. **a** Detection of firefly luciferase and GAPDH in protein lysates from EwS and non-EwS cell lines transduced with *pLenti_25_LT_Puro* by Western blot. **b** Bioluminescence measurements (exposure time: 2 min) of NSG mice 14 d after intraperitoneal injection of 1×10^7^ TU of VSV-G-pseudotyped *pLenti_25_LT* or *pLenti_CMV_LG* lentiviral particles. **c** Resazurin-based cell viability assay of *pLenti_25_LT_Puro*-transduced and selected EwS and non-EwS cell lines 72 h after GCV addition. Dots indicate relative fluorescence units normalized to vehicle control for 4 biologically independent experiments. Lines show dose-response curves with 95% confidence interval based on a three-parameter log-logistic regression model calculated for EwS or non-EwS cells respectively. **d** Annexin V/PI-staining of *pLenti_25_LT_Puro*-transduced and selected EwS and non-EwS cell lines 72 h after GCV addition. Apoptotic cells were identified as Annexin V (APC) positive cells. Dots indicate the percentage of apoptotic cells for 4 biologically independent experiments. Horizontal bars indicate mean and whiskers the standard deviation. **e** Tumor volumes of *pLenti_25_LT_Puro* pre-transduced subcutaneous xenografts. Valganciclovir (0.5 mg/ml in drinking water enriched with 5% sucrose) or sucrose (5% in drinking water) was administered orally *ad libidum* once the tumor had reached an average diameter of 5 mm. **f** Protein concentrations in conditioned medium of *pLenti_25_IX_Puro*-transduced cell lines measured by ELISA. Dots indicate calculated protein concentration for 4 biologically independent experiments. Horizontal bars indicate mean and whiskers the standard deviation for EwS or non-EwS cell lines. Concentrations below the range of detectability are not depicted in the graph. **g** Transwell Migration Assay using conditioned medium of *pLenti_25_IX_Puro*-transduced and wildtype (wt) cell lines. Migrated CD3^+^ T cells were identified and counted by flow cytometry after 4 h of incubation. Dots indicate the number of migrated CD3^+^ T cells normalized to that in the wt control for each cell line for 4 biologically independent experiments. Horizontal bars indicate mean and whiskers the standard deviation. P values were determined with two-tailed Mann-Whitney test, ****: p <= 0.0001.

These results predicted that EwS cells transduced with this vector should react with increased sensitivity to treatment with ganciclovir (GCV) compared to non-EwS cells. Indeed, when assessing cell viability after GCV treatment in resazurin-based viability assays, EwS cell lines transduced with *pLenti_25_LT_Puro* showed ∼100 fold lower effective dose 50 (ED50) concentrations than control cell lines (**Fig. 2c**). To correct for transgene independent differences in GCV-sensitivity, we also included *pLenti_0_LT_Puro-*transduced cell lines. Notably, GCV-toxicity was only induced in EwS cell lines, whereas control cell lines showed similar ED50 values for both vectors (**Additional Fig. 2c)**. In line with these observations, GCV-treatment using the average ED50 values of EwS cell lines (0.4 µM) induced extensive cell death in EwS cells but not in non-EwS controls as evidenced by Annexin V/Propidium Iodide staining and flow cytometric analysis (**Fig. 2d**). Strikingly, upon systemic treatment with Valganciclovir (VGCV) *per os*, complete tumor regression was observed in a pre-transduced EwS xenograft model (RD-ES) (**Fig. 2e**), without any detectable adverse effects, such as differences in body weight (**Addition Fig 2d**) or histomorphological changes in inner organs (not shown). Taken together, these *in vitro* and *in vivo* data, generated in EwS models, suggested that the DNA-binding preferences mediated by neomorphic functions of fusion transcription factors could be exploited to deliver a therapeutic payload with high specificity and fidelity.

To demonstrate the versatility of our expression cassette for different therapeutic approaches, we explored its suitability for tumor-specific overexpression of cytokines that may sensitize EwS for immunotherapeutic strategies, such as chimeric antigen receptor (CAR) T cell therapy. Thus, we replaced the *HSV-TK* coupled to firefly *luciferase* in *pLenti_25_LT_Puro* by the cytokines *IL-15* and *XCL1* again coupled by a P2A-linker peptide (*pLenti_25_IX_Puro*). Both cytokines are known to confer a strong activating (IL-15) and chemoattractive (XCL1) effect on T cells^43–45^. Similar to our findings with *HSV-TK*, ELISA demonstrated that EwS cells but not non-EwS control cells transduced with this new vector secreted these cytokines at relevant levels (**Fig. 2f**). Consistently, conditioned medium of EwS cells transduced with *pLenti_25_IX_Puro* was able to stimulate the migratory activity of T cells (**Fig. 2g**). Taken together, these *in vitro* data suggested that our expression cassette can be used as a flexible tool for EwS-specific expression of therapeutically exploitable genes.

### GPR64 is a promising target for targeted gene delivery in EwS

Having successfully designed and characterized a highly specific expression cassette, we sought to develop a suitable delivery strategy for therapeutic purposes *in vivo.* To increase the specificity and to enhance the viral load reaching the tumor in a therapeutic setting, we sought to combine the EwS-specific expression system with a EwS-specific transduction method, which should greatly diminish the amount of vector being lost by transducing non-target cells. Pseudotyping lentiviral particles with a modified and optimized Sindbis glycoprotein (*2.2*) has been shown to allow antibody-mediated transduction *in vivo*^46,47^. While in principle CD99 would constitute a highly expressed surface protein in EwS, its ubiquitous expression in normal tissues renders this protein unsuitable for such an approach^32^. To identify EwS-specific candidate surface proteins that are highly expressed in EwS but only minimally in normal tissues, we analyzed a previously described set of gene expression microarray data from 50 EwS and 928 normal tissues (comprising 70 tissue types) and identified 36 genes that were significantly overexpressed in EwS compared to any other normal tissue (**Additional Table 2**)^32^. Of these, 3 genes (*GPR64*, *FAT4* and *LECT1*) encoding cell surface proteins were selected for *in vitro* analysis based on the availability of commercial monoclonal antibodies targeting their extracellular domains (**Fig. 3a, Additional Fig. 3**). Indirect antibody staining and flow cytometry analysis confirmed the surface-expression of GPR64 and, to a lesser extent, of FAT4 in 6 EwS cell lines at the protein level (**Fig. 3b**). Interestingly, the membrane-bound disialoganglioside GD2, which was recently identified as potential target for antibody- or CAR T cell-based therapies, showed only a weak staining signal in the 6 EwS cell lines tested^48^. Thus, due to its higher expression levels, GPR64 was selected for further experiments and its specific expression was confirmed *in situ* in patient-derived EwS tumor tissue (n = 18) and normal tissues (n = 29) by immunohistochemistry (**Fig. 3c**). Notably, apart from the epididymis, only minimal GPR64 expression was found in any other organ whereas the majority of EwS samples showed positive staining in immunohistochemistry (**Additional Table 3**).

**Fig. 3:**
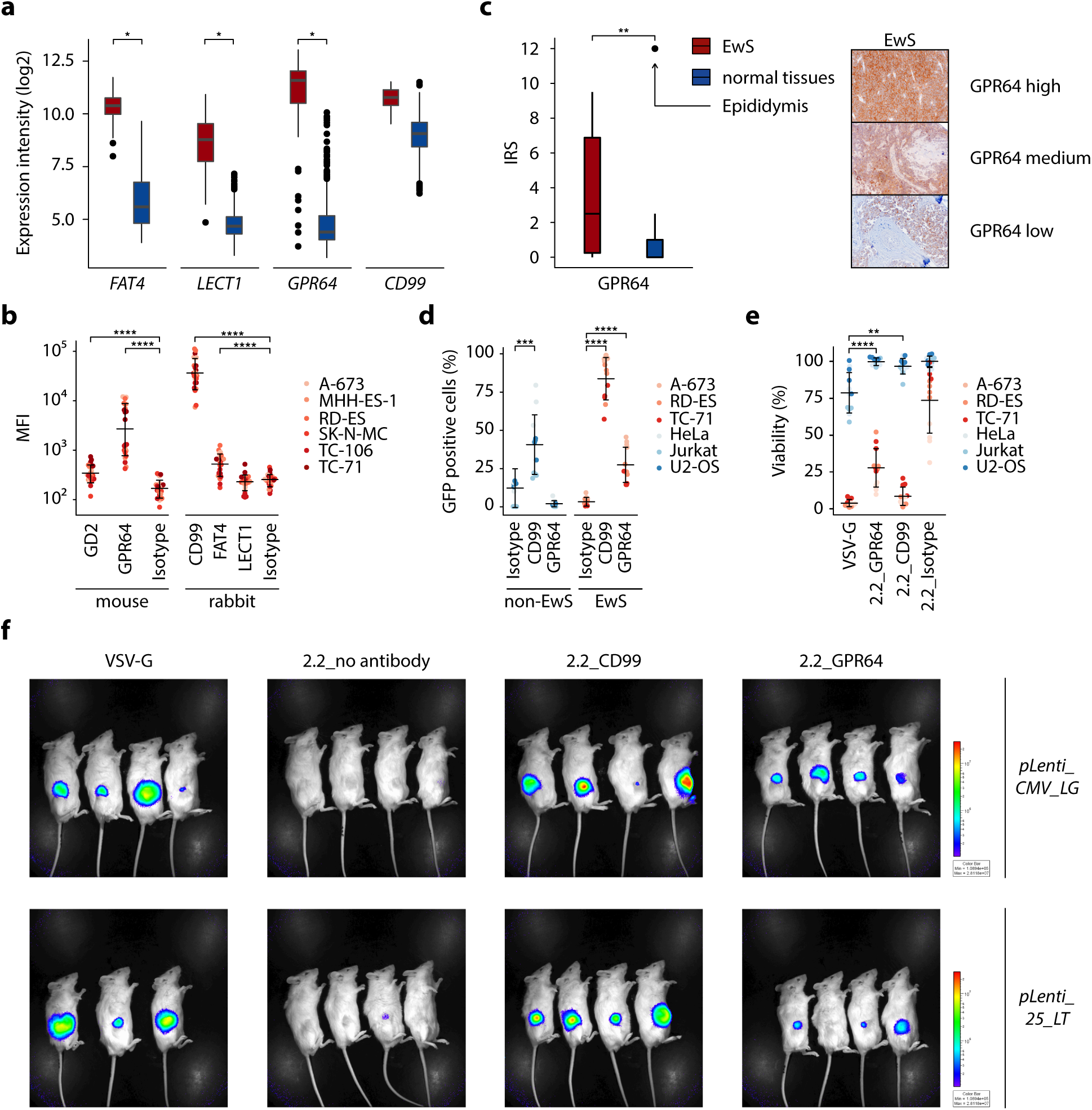
GPR64 allows EwS-specific lentiviral transduction. **a** mRNA log2 expression intensities of *GPR64*, *FAT4*, *LECT1*, and *CD99* from publicly available microarray data of EwS (n = 50) and normal tissues (n = 928, comprising 70 different tissue types). Data are presented as boxplots with the horizontal line representing the median, the box the interquartile range (IQR) and the whiskers 1.5 * IQR of the expression intensity. **b** Validation of surface expression of GD2, GPR64, CD99, FAT4 and LECT1 by antibody staining and flow cytometry. Isotype controls for both antibody host species were included separately. Dots indicate mean fluorescent intensity (MFI) for 4 independent experiments. Mean and standard deviation per group are depicted as horizontal bars and whiskers. **c** IRS (immunoreactive score) of GPR64 in immunohistochemistry of primary EwS tumors and relevant normal tissues. Representative EwS samples with high, medium and low GPR64 expression are shown aside. **d** Flow cytometry analysis of EwS and non-EwS cell lines after transduction with GPR64-targeting, *GFP*-encoding lentiviruses. CD99 and isotype-targeting lentivirus was used as positive and negative control. Dots indicate percentage of GFP positive cells determined by flow cytometry of 4 biologically independent experiments. Horizontal bars and whiskers represent mean and standard deviation per group. **e** Resazurin-based cell viability assay of EwS and non-EwS cell lines treated with GCV (20 µM) or DMSO vehicle control 24h after GPR64-targeted transduction with *pLenti_25_LT.* Readout was performed 72h after GCV addition. CD99-targeting lentiviruses, non-targeting lentiviruses (isotype) and VSV-G pseudotyped lentiviruses were included as controls. Dots indicate cell viability relative to that of vehicle control for 4 biologically independent experiments. Mean standard deviation per group are represented by horizontal bars and whiskers. **f** Bioluminescence measurements (exposure time: 20 sec) of NSG mice bearing subcutaneous RD-ES xenografts 14 d after a single intratumoral injection of 0.5 × 10^6^ TU of *pLenti_25_LT* or *pLenti_CMV_LG* lentiviral particles pseudotyped with 2.2. GPR64- or CD99-targeting antibodies were used to coat 2.2 pseudotyped viruses. VSV-G pseudotyped lentiviruses were used as positive control, 2.2 pseudotyped viruses without antibodies were included as negative control. P values were determined with two-tailed Mann-Whitney test, * :p <= 0.05, **: p <= 0.01, ***: p <= 0.001, ****: p <= 0.0001.

To evaluate the suitability of GPR64 as a candidate for targeted transduction of EwS cells, lentiviral particles were produced using a transfer plasmid containing a GFP reporter expressed by a CMV promoter and the 2.2 packaging plasmid. Next, EwS (A-673, RD-ES, TC-71) cell lines and non-EwS (HeLa, Jurkat, U2-OS) control cell lines were transduced with these vectors combined with either a GPR64 antibody, a CD99 antibody, or an isotype control. Remarkably, flow cytometry analysis showed specific GFP expression of EwS cells when targeting GPR64 while no significant GFP-positivity was seen in control cells or when isotype-coated virus was added (**Fig. 3d**). In accordance with its ubiquitous expression, CD99-coated viral particles showed non-specific transduction of both EwS and control cell lines.

To assess whether the addition of this transduction-based targeting strategy could further increase the therapeutic specificity of our transcription-based approach, 3 EwS and 3 control cell lines were treated with equal amounts of lentivirus either pseudotyped by VSV-G, or antibody-coated 2.2 (GPR64, CD99 or isotype control) using the aforementioned transfer plasmid *pLenti_25_LT*. Subsequent addition of GCV (20 µM) revealed a significant reduction in GCV sensitivity in non-EwS control cells treated with GPR64-coated viral particles compared to those treated with VSV-G pseudotyped virus, underlining the benefit of the additional EwS-specific delivery strategy (**Fig. 3e**).

Next, we aimed to investigate whether antibody-mediated transduction of EwS cells was also feasible *in vivo*. To this end, we intratumorally injected either VSV-G- or 2.2-pseudotyped and antibody-coated lentiviral particles carrying the firefly *luciferase* transgene under control of our EwS-specific expression cassette (*pLenti_25_LT*) or driven by the ubiquitous CMV promoter (*pLenti_CMV_LG*) (**Fig. 3f**). Notably, in RD-ES xenografts, comparable tumor-derived luminescence was detected when injecting GPR64- or CD99-directed, 2.2-pseudotyped compared to VSV-G-pseudotyped lentiviruses that served as positive control. Moreover, plain, uncoated 2.2-pseudotyped viruses achieved no detectable transduction both in the *pLenti_25_LT* and *pLenti_CMV_LG* group. These results were confirmed in a second cell line (A-673) (**Additional Fig. 4**). In sum, these experiments demonstrate the feasibility of antibody-mediated targeted transduction of EwS cells *in vivo*.

### The combination of EwS-specific delivery and gene expression improves specific tumor therapy *in vivo*

Having established both, a EwS-specific expression cassette and delivery strategy, we moved on to combine these two for therapeutic purposes *in vivo*. Therefore, we subcutaneously inoculated A-673 EwS cells and, once the average tumor diameter had reached 5 mm, intratumorally injected GPR64-directed, 2.2-pseudotyped treatment (*pLenti_25_LT*) or mock (*pLenti_CMV_LG*) virus. Excitingly, upon oral VGCV administration, most tumors in the *pLenti_25_LT*-transduced group showed significant reduction in tumor growth compared to the control groups (**Fig. 4a**). In a second step, we evaluated the efficacy of our treatment strategy in a more systematic setting by inoculating luciferase-expressing A-673 cells intraperitoneally and repeatedly injecting GPR64-directed, 2.2-pseudotyped lentivirus expressing *HSV-TK* (*pLenti_25_TK*) or PBS (negative control) into the peritoneal cavity 3 days after tumor inoculation. Excitingly, while the control group showed a strong increase of luminescence over time corresponding to strong increase in peritoneal tumor mass, a significantly lower bioluminescent signal was detected in the treatment group (**Fig. 4b** and **Additional Fig. 5**).

**Fig. 4:**
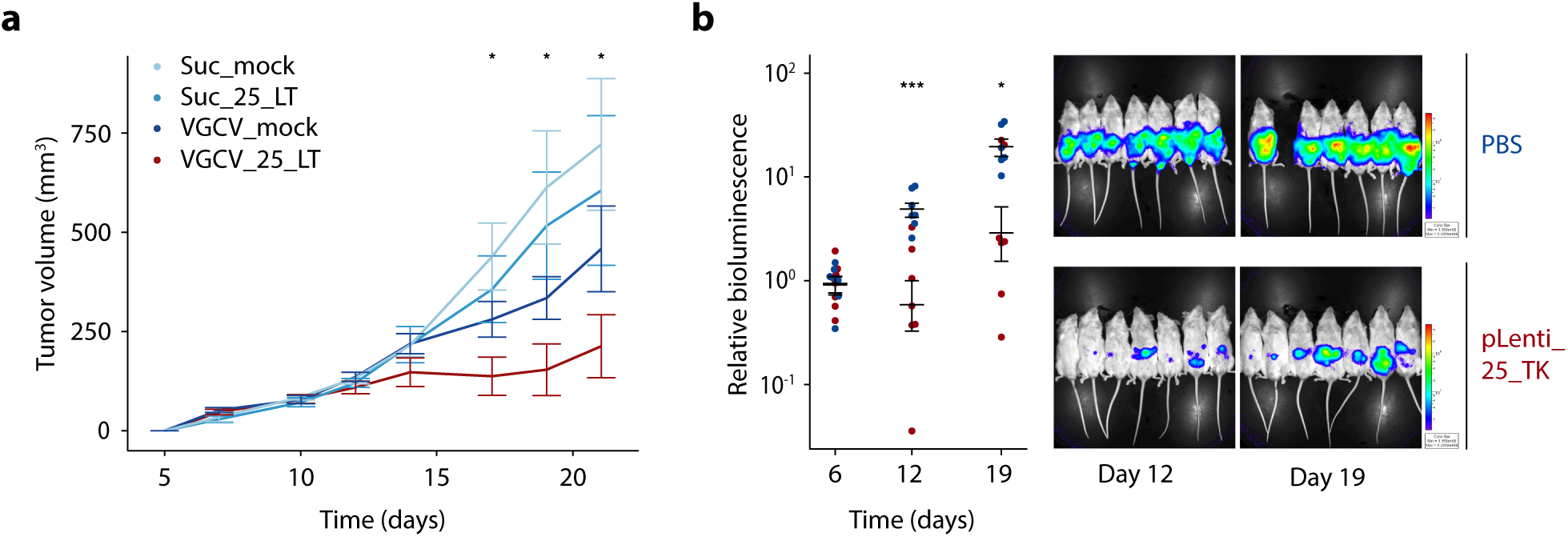
The combination of EwS-specific gene expression and delivery enables effective tumor therapy *in vivo*. **a** Tumor volumes of A-673 subcutaneous xenografts treated with GPR64-targeting *pLenti_25_LT* or *pLenti_CMV_LG* (mock) lentiviruses. Valganciclovir (VGCV, 0.5 mg/ml in drinking water enriched with 5% sucrose) or sucrose (5% in drinking water) was administered orally *ad libidum* once the tumor had reached an average diameter of 5 mm. Lentiviruses were intratumorally injected twice per week starting from day 7. Data are shown as mean tumor volume and SEM of 6-7 mice per treatment condition. P values were determined by one-tailed Mann-Whitney test. **b** Relative bioluminescence (right) and bioluminescent images (left) of NSG mice after intraperitoneal tumor inoculation with firefly luciferase-expressing A-673. 3 days after tumor injection mice were randomized and repeatedly received either GPR64-directed 2.2. pseudotyped lentivirus (*pLenti_25_TK*) or PBS by intraperitoneal injection. VGCV was orally administered in both groups 3 days after the first virus injection. The representative bioluminescent pictures show both groups 12 and 19 days after tumor inoculation. Dots indicate bioluminescence signal relative the mean measured on of VGCV initiation (day 6) for 6-7 mice per group. Horizontal bars indicate mean and whiskers SEM per group. P values were determined by one-tailed Mann-Whitney test. * :p <= 0.05, ***: p <= 0.001.

In conclusion, our results indicate that the neomorphic aberrant DNA-binding properties of EF1 enable EwS specific and EF1-dependent expression of therapeutic transgenes *in vivo*.

### Highly specific, enhancer-based gene expression systems can be designed for other fusion-driven pediatric sarcomas

To investigate whether this principle can be translated to other cancers driven by an oncogenic fusion transcription factor, we extended our analyses to fusion-positive ARMS, which harbors the dominant chimeric P3F1 oncoprotein in more than 50% of cases^49^. P3F1 mediates cell transformation by binding to specific DNA motifs thereby establishing *de novo* super-enhancers (SEs) encompassing known oncogenes, such as *ALK*, causing dysregulating of the transcriptome^5,8,50^. Thus, we first cloned a ∼300 bp DNA segment (chr2:29,657,671–29,657,976; hg38) from the third intron of *ALK*, that has been identified as a strong P3F1-binding site, into a luciferase reporter plasmid upstream of YB-TATA^5^. Similar to our observations made in EwS (**Fig. 1c**), we found a significant induction of reporter gene expression in fusion-positive ARMS (RH4 and RH30) but not in fusion-negative embryonal rhabdomyosarcoma (RD) or in non-rhabdomyosarcoma control cell lines (U2-OS, HeLa, Jurkat and A-673) (**Fig. 5a**). Interestingly, Gryder et al. showed that two point mutations of a single P3F1-binding motif, consisting of GTCACGGT, abrogated the transactivating activity of the ALK-SE^5^. To further improve the induction capacity of this construct, we optimized the sequence at this putative P3F1 binding site to completely match the ATTW**GTCACGGT** motif (syn_alk) as annotated by HOMER motifs, which resulted in improved luciferase signals in fusion positive ARMS cell lines but not in control cell lines (**Fig. 5b**). Strikingly, the fusion positive ARMS-specific expression induction could be further increased by adding three (syn_alk_3) or five (syn_alk_5) additional ATTW**GTCACGGT** motifs to the SE sequence (**Fig. 5b**).

**Fig. 5:**
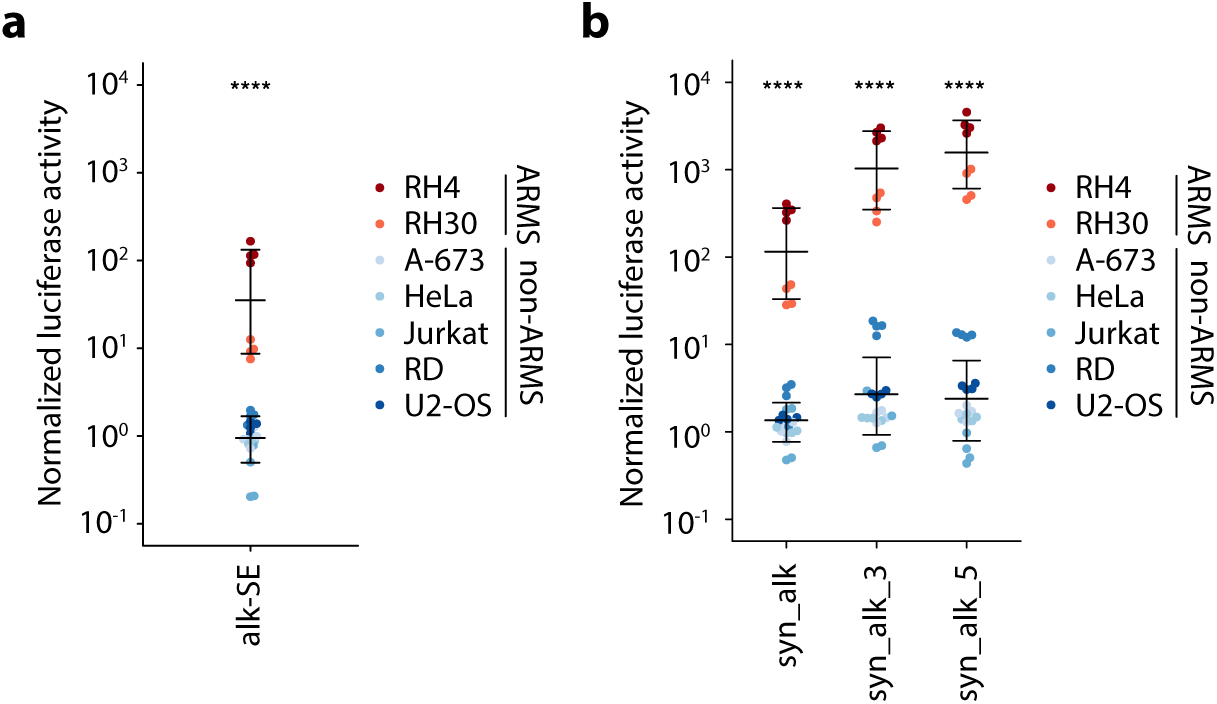
Highly specific, enhancer-based gene expression systems can be designed for other fusion-driven pediatric sarcomas. **a** Luciferase reporter assays of indicated fusion-positive ARMS (RH4 and RH30) and control cell lines after co-transfection with a reporter plasmid containing the alk-SE upstream of the minimal promoter YB-TATA and a constitutively expressed *Renilla* luciferase-encoding plasmid. Dots indicate firefly to *Renilla* luminescence ratios normalized to a reporter plasmid without the alk-SE. Horizontal bars indicate mean and whiskers standard deviation per group. **b** Luciferase reporter assays of the same cell lines as in Fig. 5a after co-transfection with a reporter plasmid containing the either syn_alk, syn_alk_3 or syn_alk_5 upstream of the minimal promoter YB-TATA and a constitutively expressed *Renilla* luciferase-encoding plasmid. Dots indicate firefly to *Renilla* luminescence ratios normalized to a reporter plasmid without the alk-SE. Horizontal bars indicate mean and whiskers standard deviation per group. P values were determined by two-tailed Mann-Whitney test, ****: p <= 0.0001.

Collectively, these results indicate that our approach is translatable to other cancers driven by oncogenic transcription factors with unique DNA-binding properties.

## Discussion

Increasingly standardized classification systems and risk-stratification-based multimodal therapy have strongly improved outcomes of patients affected by pediatric sarcomas including EwS and ARMS, especially in localized disease^51,52^. However, patients with relapsed or recurrent or metastatic disease still have dismal prognosis despite the often mutilating and highly toxic nature of the applied therapies^1,53^.In addition, only a few targeted therapies have been tested in early phase clinical trials yet and showed limited or no activity in both entities^54,55^. Thus, innovative treatment strategies are urgently needed to not only improve the prognosis of affected children and adolescents but also reduce the side effects of standard therapy.

Although EF1 and P3F1 would represent ideal therapeutic targets, oncogenic fusion transcription factors typically lack enzymatic activity and are as such often considered to be undruggable^56^. Importantly, accumulating evidence in EwS suggests that low levels of EF1 expression (aka activity) may rather foster metastasis than acting tumor suppressive, possibly rendering approaches that aim at therapeutically lowering the fusion oncogenes activity futile^57–60^. In stark contrast, our approach aims at exploiting the neomorphic DNA-binding properties of chimeric fusion-oncoproteins for tumor-specific therapeutic gene expression.

Gene therapy and gene editing have been rapidly evolving in cancer over the past decades and might soon expand the armory of cancer therapeutics in clinical use^61–64^. However, compared to the recent success of gene therapy in monogenic hereditary diseases such as hemophilia, cancer gene therapy lags behind due to difficulties regarding safety and delivery^65^. To improve the safety of the genetic payload, tumor-specific gene expression cassettes have been generated for a multitude of malignancies with varying success^66,67^. Whereas most of these designs focused on endogenous promoter sequences that are targeted by transcription factors over-expressed in the respective cancer entity^68–70^, to the best of our knowledge no pure enhancer-based design has been reported so far. Moreover, we are not aware of a single study that exploited neomorphic, and hence unique, binding properties of tumor-defining driver oncoproteins. In contrast to endogenous promoter sequences that enable gene expression in any cell that expresses the required *wildtype* transcription factors, our design harnesses *de-novo* DNA binding of pathognomonic *mutated* oncoproteins that do not exist in non-transformed, healthy tissue. Further, we proved the versatility of our design by using several transgenes (*gfp*, *luciferase*, *HSV TK SR-39*, *IL-15* and *XCL1*) and observed functional gene expression levels both *in vivo* and *in vitro* in two different sarcoma entities (EwS and ARMS). Moreover, we showed the safety and effectiveness of entirely synthetic cassettes lacking genomic flanking regions of the regulatory core elements (i.e. GGAA-msats) in preclinical models. Thus, our design might serve as a blueprint for similar expression systems across a wide variety of cancer entities in the future.

As mentioned above, transgene or Cas9-ribonucleoprotein delivery has been a major bottleneck in gene therapy and gene editing. For our study, we chose the well characterized lentiviral delivery system, as it has been successfully employed for *in vivo* transgene delivery in the past^71^. Moreover, lentiviruses allow vector pseudotyping and thereby represent a versatile system for cell- or tissue-specific gene delivery^46^. Performing *in silico* analyses and *in vitro* experiments, we identified GPR64 as a promising surface antigen for EwS-specific gene delivery by antibody-mediated transduction. GPR64 has been previously shown to be expressed in EwS and was associated with increased invasiveness and metastasis, possibly rendering GPR64 as an especially relevant target structure in metastatic disease^72^. Despite its lower expression levels compared to CD99, GPR64-targeted transduction was highly EwS-specific and allowed sufficient gene transfer *in vivo* after intratumoral and intraperitoneal injection.

The therapeutic targeting approach presented in this study is unprecedented in the field of fusion-gene driven pediatric sarcoma. The design of our synthetic expression cassette exploits the strong oncogene-dependence of EwS and ARMS, and harnesses unique, neomorphic DNA-binding properties of their pathognomonic driver-oncogenes. However, our strategy faces the same difficulties and obstacles as other gene editing or gene delivery approaches in cancer therapy.

For its strong and well-characterized phenotype, we chose *HSV-TK* as transgene for most of our experiments. Despite our and others’ promising results in preclinical models, *suicide gene therapy* is limited to localized disease in clinical use as high transduction efficiencies are often not achievable upon systemic administration^62^. Thus, other transgenes, i.e. immunostimulatory cytokines, might be a better therapeutic payload as they do not require high transduction efficiencies to exert their function in combination with other immunotherapeutic strategies, such as CAR T cell therapies. Of note, and as for most other solid tumors, no effective CAR T cell therapy is currently available for EwS and ARMS, both of which represent immunologically rather ‘cold’ tumors^73^. As a proof-of-concept, we therefore chose *IL-15* and *XCL1* as suitable transgenes for immunostimulatory priming of the tumor microenvironment.

IL-15 shows similarities to IL-2 and was found to regulate survival and activation of T cells. Moreover, it was found to enhance *in vivo* antitumor activity of CD8^+^ T cells and has been included in CAR designs as tethered cytokine ^43,44^. The chemokine XCL1 was shown to be mainly expressed and secreted in activated CD8^+^ T cells and to be crucial for effective antigen cross-presentation by dendritic cells (DC) and antigen spreading of tumor infiltrating lymphocytes^45,74^. Whereas XCR1, the receptor for XCL1, seems to be exclusively expressed in CD8^+^ DC in mice, in humans XCR1 expression was also detected in a wide range of T cells and natural killer (NK) cells apart from myeloid DCs^45,75^. Indeed, XCL1 was shown to act as a chemoattractant on human T cells in several studies^76,77^. The combination of both transgenes is a promising approach to prime the tumor microenvironment for successful CAR T cell therapies in pediatric sarcoma and we are keen to evaluate its effect *in vivo* in future studies, especially as EwS-targeting CAR or TCR-transgenic T cells have been previously generated^78,79^.

Moreover, integrating viral vectors, such as lentiviruses, will ultimately be replaced by non-integrating, episomal vectors such as Adeno-associated Viruses (AAV) due to their better safety profile. Although restricting the tropism of non-lentiviral vectors to one tissue or cell type is much more difficult than in lentiviruses, receptor targeting AAV vectors have been generated in the past^80^. Notably, our study showed the functionality and specificity of our design even when delivered episomally, thus allowing an easy transfer to episomal viral delivery systems once they are available for pediatric sarcoma.

## Conclusion

In summary, our results provide evidence that the unique interaction of oncogenic fusion transcription factors with aberrant binding sites can be used for specific therapeutic gene expression. In this study, we demonstrate this using a broad range of transgenes in EwS and fusion-positive ARMS. Besides the often poor specificity of ‘cancer-specific’ expression systems **–** a problem that we successfully addressed in this study **–** the limited success in delivering transgenes to a sufficient fraction of tumor cells has been the main obstacle of cancer gene therapy. Hence, it is of utmost importance for gene delivery strategies to be refined and improved for future translation of innovative and specific cancer gene therapies into the clinics.

As the fusion oncogene-based expression systems proposed by us are of a simple architecture and show tumor specificity both in integrating as well as episomal vectors, they could be used for any therapeutic approach relying on transgene expression in cancer cells, including replicating oncolytic viruses.

## List of abbreviations

APC: Allophycocyanin
ARMS: Alveolar rhabdomyosarcoma
CMV: Cytomegalovirus
DMSO: Dimethyl sulfoxide
Dox: Doxycycline
ED: Effective dose
EF1: EWSR1-FLI1
EwS: Ewing sarcoma
FITC: Fluorescein
GCV: Ganciclovir
GFP: Green fluorescent protein
HSV: Herpes simplex virus
IRS: Immunoreactive score
KD: Knockdown
LG: Luciferase-P2A-GFP
LT: Luciferase-P2A-thymidine kinase
MFI: Mean fluorescence intensity
P3F1: PAX3-FOXO1
PE: Phycoerythrin
PI: Propidium iodide
Puro: Puromycin
RT: Room temperature
SE: Super-enhancer
TK: Thymidine kinase
TU: Transducing units
WT: Wildtype

## Declarations

### Ethics approval and consent to participate

Human tissue samples were retrieved from the tissue archives of the Institute of Pathology of the LMU Munich (Germany) upon approval of the institutional review board. All patients provided informed consent. Tissue-microarrays (TMAs) were stained and analyzed with approval of the ethics committee of the LMU Munich (approval no. 307–16 UE).

### Availability of data and materials

All ChIP-seq, RNA-seq and Microarray data reanalyzed as part of this study are publicly available under the accession codes listed in Additional Table 5 and the methods section.

### Authors’ contributions

T.L.B.H., T.G.P.G. and M.M.L.K. conceived the study. T.L.B.H., T.G.P.G. and M.M.L.K. wrote the paper, and drafted the figures and tables. T.L.B.H., M.M.L.K., I.P. and J.L. carried out *in vitro* experiments. T.L.B.H. performed bioinformatic and statistical analyses. T.L.B.H., M.M.L.K., D.M., B.P., S.J., S.O. and F.C.A. performed and/or coordinated *in vivo* experiments. S.L. performed immunoreactive scoring. B.L.C., S.O., S.J.J. and J.Z. contributed to experimental procedures. A.K. provided excellence guidance with T cell engineering and T cell transfer. D.A., and F.K. provided laboratory infrastructure.

## Acknowledgements

We wish to thank Prof. Thomas Kirchner for providing lab space and scientific support.

Moreover, we thank Laura Hippe, Stefanie Stein, Anja Heier and Andrea Sendelhofert for expert technical assistance and Benjamin Knust for his assistance with animal experiments. We thank the CellSort Team of TU Munich for assistance with flow cytometry analysis and cell sorting.

## Additional figure legends

**Additional Fig. 1**

**a** Epigenetic profile of the *FEZF1-AS1* locus in indicated EwS cells transduced with either a control shRNA (shGFP) or a specific shRNA against *EF1* (shEF1) from published ChIP-seq data for EWSR1-FLI1 and H3K27ac^6^. **b** Volcano plot of published RNA-seq data showing differentially expressed genes (DEGs) after shRNA-mediated *EF1* (shEF1) knockdown compared to a non-targeting shRNA (shGFP). A summary of two cell lines is shown (A-673 and SK-N-MC). *FEZF1-AS1* is depicted in red. **c** Luciferase reporter assays of A-673/TR/shCtrl co-transfected with the same plasmids as in Fig. 1c treated with / without Dox. Dots indicate firefly to *Renilla* luminescence ratios normalized to a reporter plasmid without GGAA-repeats for 4 biologically independent experiments. Horizontal bars indicate mean and whiskers standard deviation per group. **d** Analysis of *EF1* mRNA expression after 72 h of doxycycline treatment compared to untreated controls in A-673/TR/shEF1 cells by RT-qPCR. Dots indicate gene expression relative to untreated controls by determined by the 2^-ΔΔCT^ method for 4 independent experiments. Horizontal bars indicate mean and whiskers standard deviation per group.

**Additional Fig. 2**

**a** Analysis of *HSV TK* mRNA expression in *pLenti_25_LT_Puro*-transduced and selected EwS and non-EwS cell lines compared to *pLenti_0_LT_Puro*-transduced and selected EwS and non-EwS cell lines. Dots indicate ΔCT values of *TK* compared to *RPLP0*. Horizontal bars indicate mean and whiskers standard deviation per group. **b** Analysis of *WPRE* copy numbers in indicated organs after intrapertioneal injection of VSV-G pseudotyped *pLenti_25_LT* or *pLenti_CMV_LG* lentiviral particles by genomic qPCR. Dots indicate ΔCT values of *WPRE* compared to murine *ACTB* for 4 mice per group. Horizontal bars indicate mean and whiskers standard deviation per group. **c** Resazurin-based cell viability assay of *pLenti_25_LT_Puro*-*pLenti_0_LT_Puro*-transduced and selected EwS and non-EwS cell lines 72 h after GCV addition. Dots indicate relative fluorescence units normalized to vehicle control for 4 biologically independent experiments. Lines show dose-response curves with 95% confidence interval based on a three-parameter log-logistic regression model calculated for EwS or non-EwS cells respectively. **d** Weight curves of VGCV-(0.5 mg/ml) treated and untreated (sucrose) tumor bearing NSG mice. Lines indicate mean weights of 7-10 mice per group and whiskers standard error of mean.

P values were determined by two-tailed Mann-Whitney test, ****: p <= 0.0001.

**Additional Fig. 3**

mRNA log2 expression intensities of **a** *FAT4*, **b** *LECT1* and **c** *GPR64* from publicly available microarray data of EwS (n = 50) and normal tissues (n = 928, comprising 70 different tissue types). Data are presented as boxplots with the horizontal line representing the median, the box the interquartile range (IQR) and the whiskers 1.5 * IQR of the expression intensity.

**Additional Fig. 4**

Bioluminescence measurements (exposure time: 3 sec) of NSG mice bearing subcutaneous A-673 xenografts 14 d after a single intratumoral injection of 0.5 × 10^6^ TU of *pLenti_25_LT* or *pLenti_CMV_LG* lentiviral particles pseudotyped with 2.2. Anti-GPR64 antibody was used to coat 2.2 pseudotyped viruses. 2.2 pseudotyped viruses without antibody were included as negative control.

**Additional Fig. 5**

Bioluminescent images of NSG mice (exposure time: 2 sec) after intraperitoneal tumor inoculation with luciferase-expressing A-673. 3 days after tumor injection mice were randomized and repeatedly received either GPR64-directed 2.2. pseudotyped lentivirus (*pLenti_25_TK*) or PBS by intraperitoneal injection. VGCV was orally administered in both groups 3 days after the first virus injection. The representative bioluminescent pictures show both groups 12 and 19 days after tumor inoculation. The time-line below depicts the detailed design of the experiment.

**Additional Fig. 6**

**a** Representative flow cytometry gating strategy for identification of apoptotic EwS and non-EwS control cell lines after GCV treatment. Dot plots show a representative sample of SK-N-MC cells treated with 0.4 µM GCV for 72 h. Annexin V was stained by APC, PI by PE. **b** Representative flow cytometry gating strategy for identification of migrated T cells and counting beads. Dot plots show a representative sample of PBMC migrated towards conditioned medium of *pLenti_25_IX_Puro* pre-transduced and selected A-673. PI was stained by PE, CD3 was stained by FITC. **c** Representative flow cytometry gating strategy for identification of surface antigen expression by indirect staining procedure. Dot plots show a representative sample of A-673 cells indirectly stained for GPR64 (APC) after exclusion of dead cells by PI (PE). **d** Representative flow cytometry gating strategy for identification of GFP-transduced EwS and non-EwS control cell lines after antibody-mediated transduction. Dot plots show a representative sample of A-673 cells transduced with GPR64-targeting vectors.

