## Supplementary figures and images for "Neomorphic DNA-binding enables tumor-specific therapeutic gene expression in fusion-addicted childhood sarcoma"

### Additional Figures

**Additional Fig. 1**

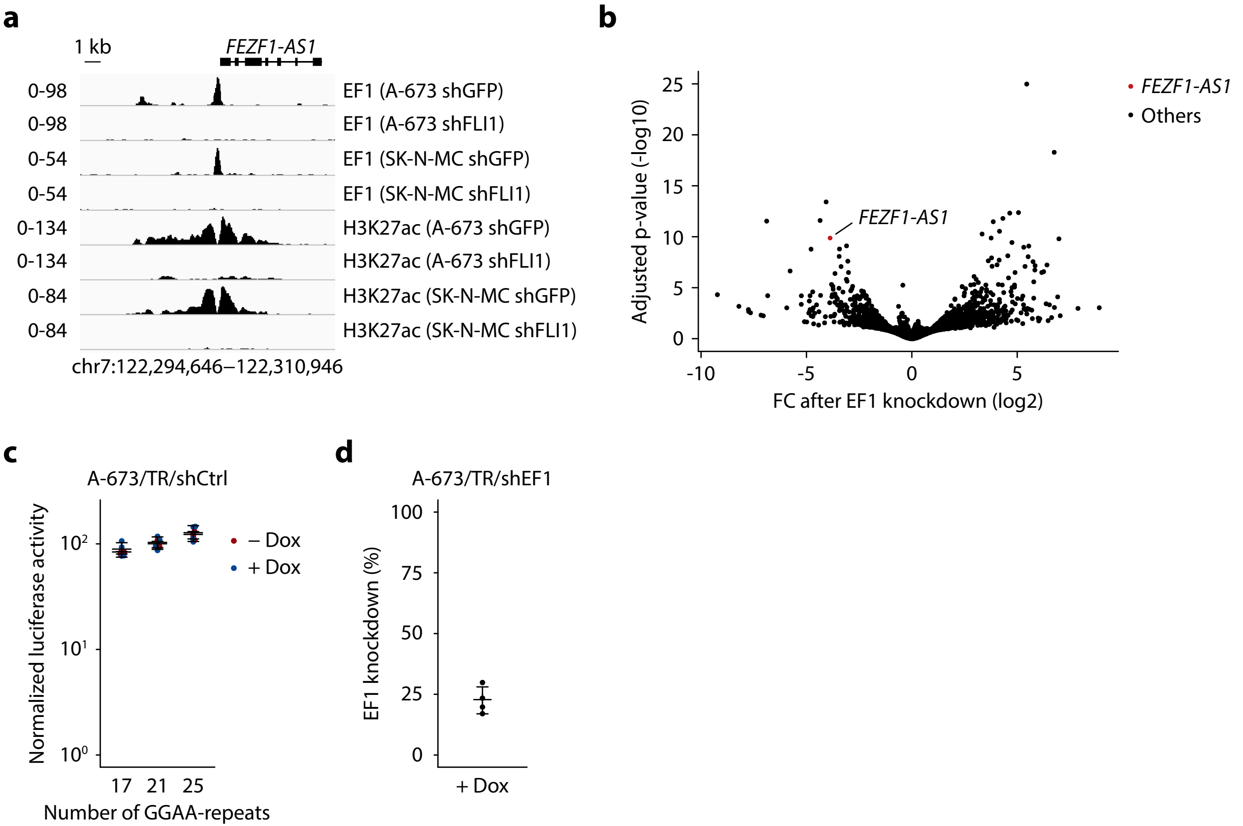

##### Additional Fig. 2

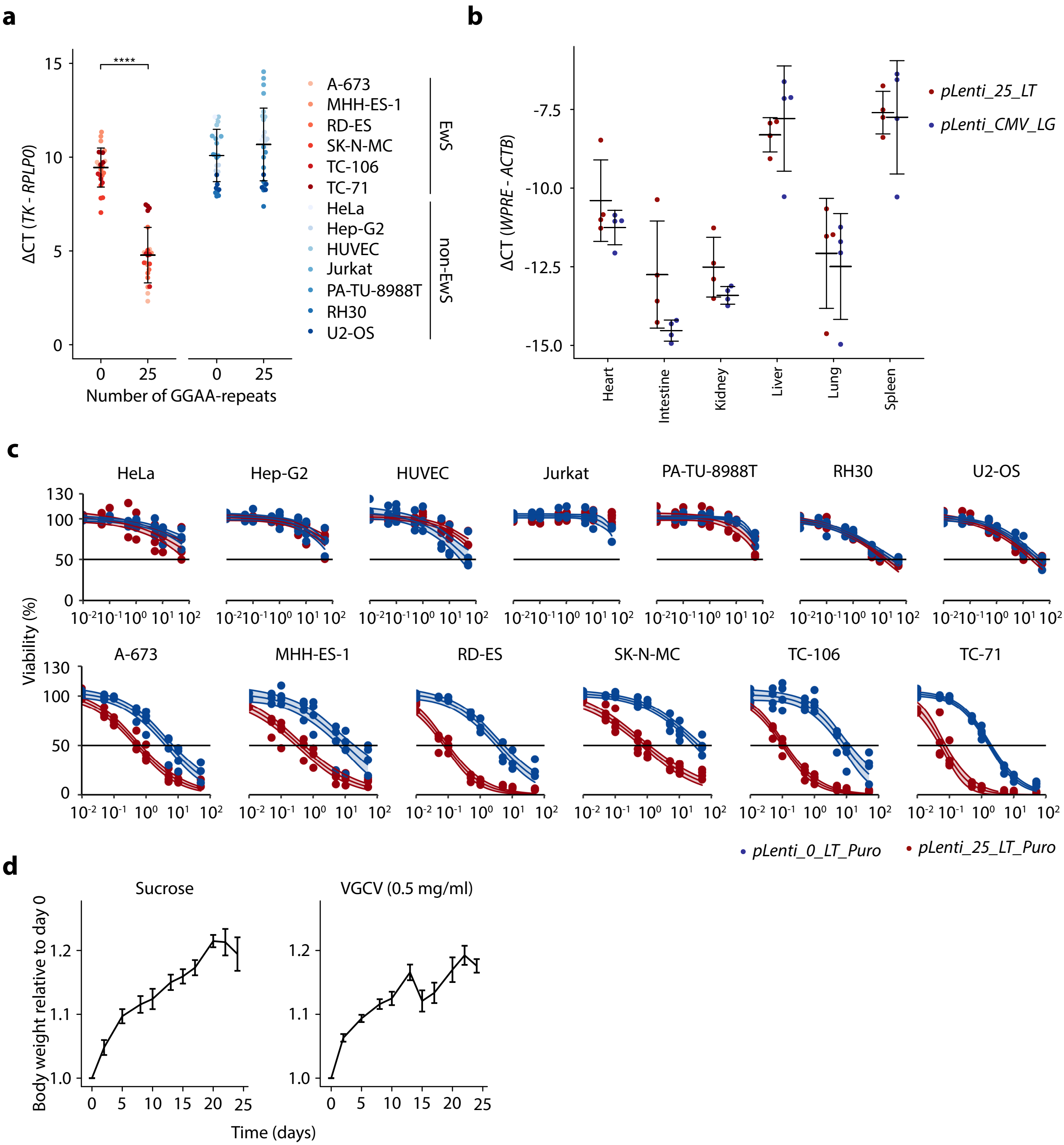

##### Additional Fig. 3

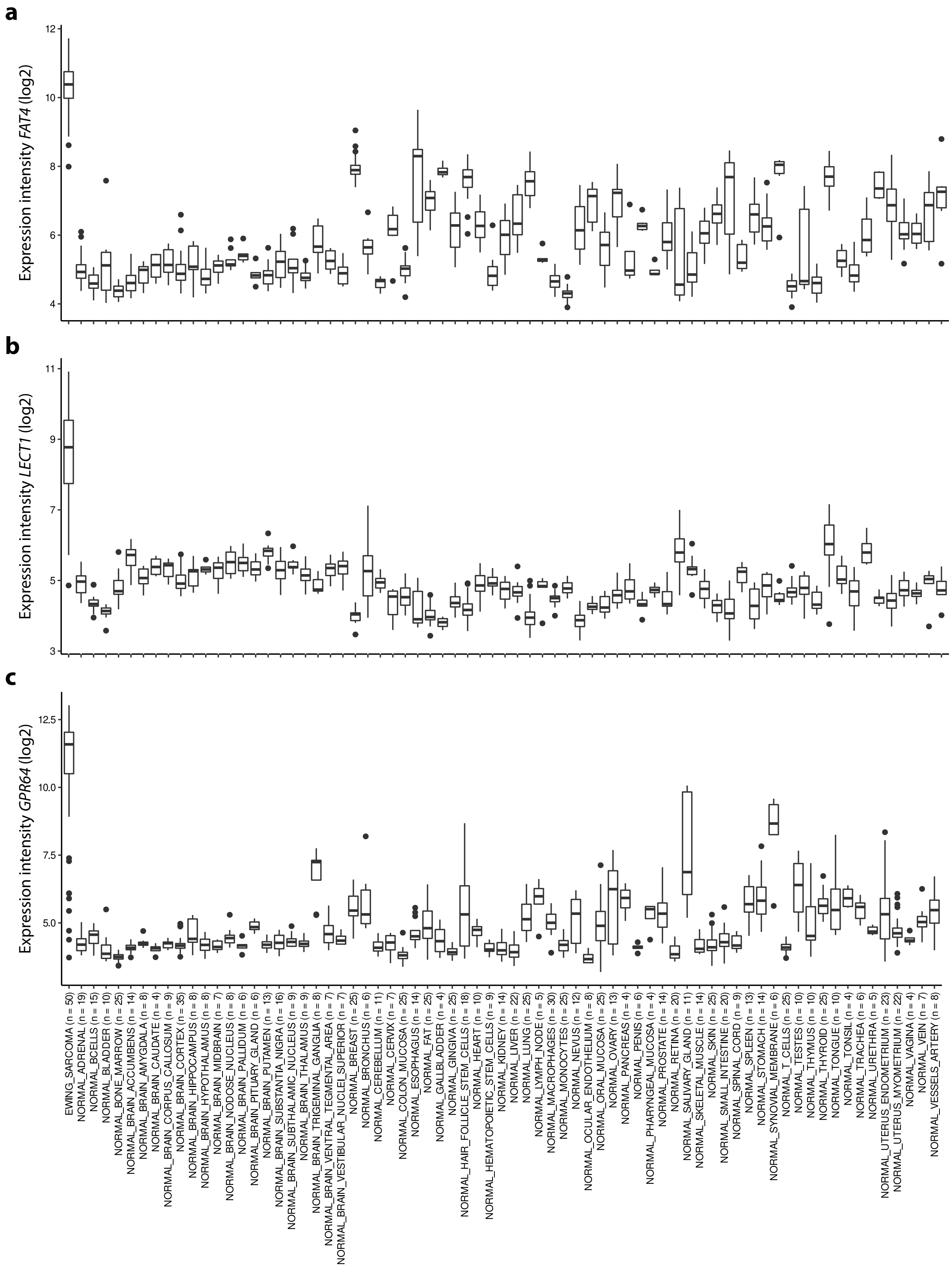

#### Additional Fig. 4

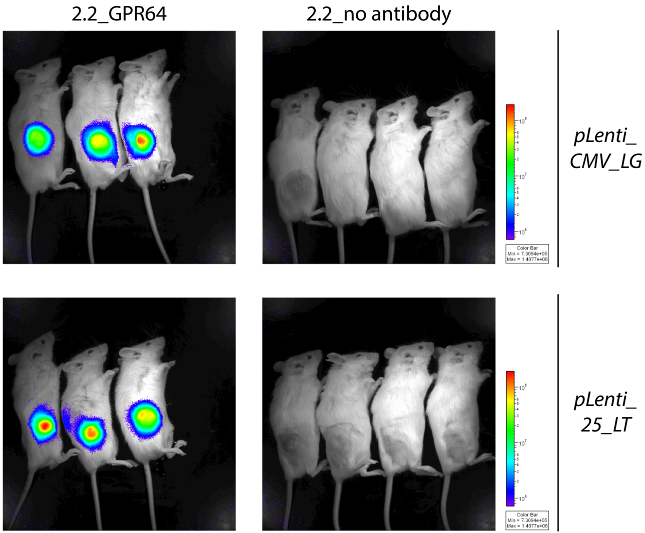

#### Additional Fig. 5

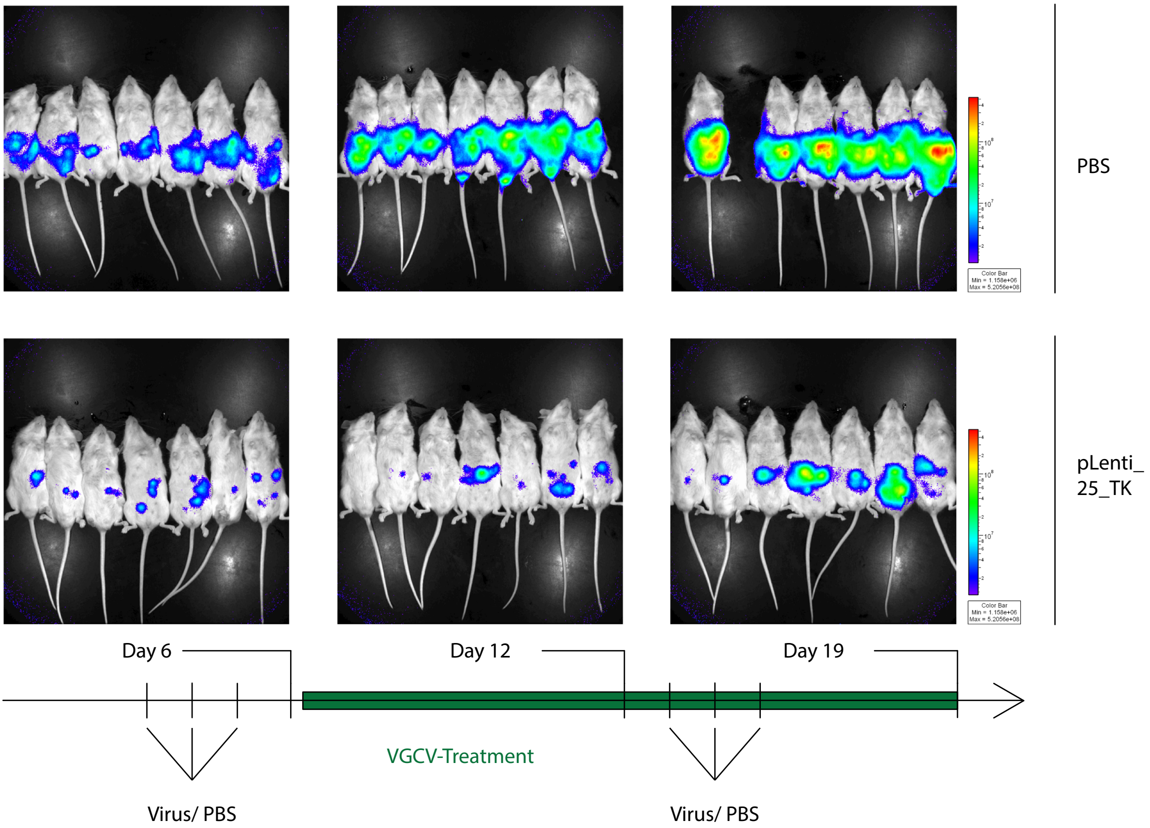

**Additional Fig. 6**

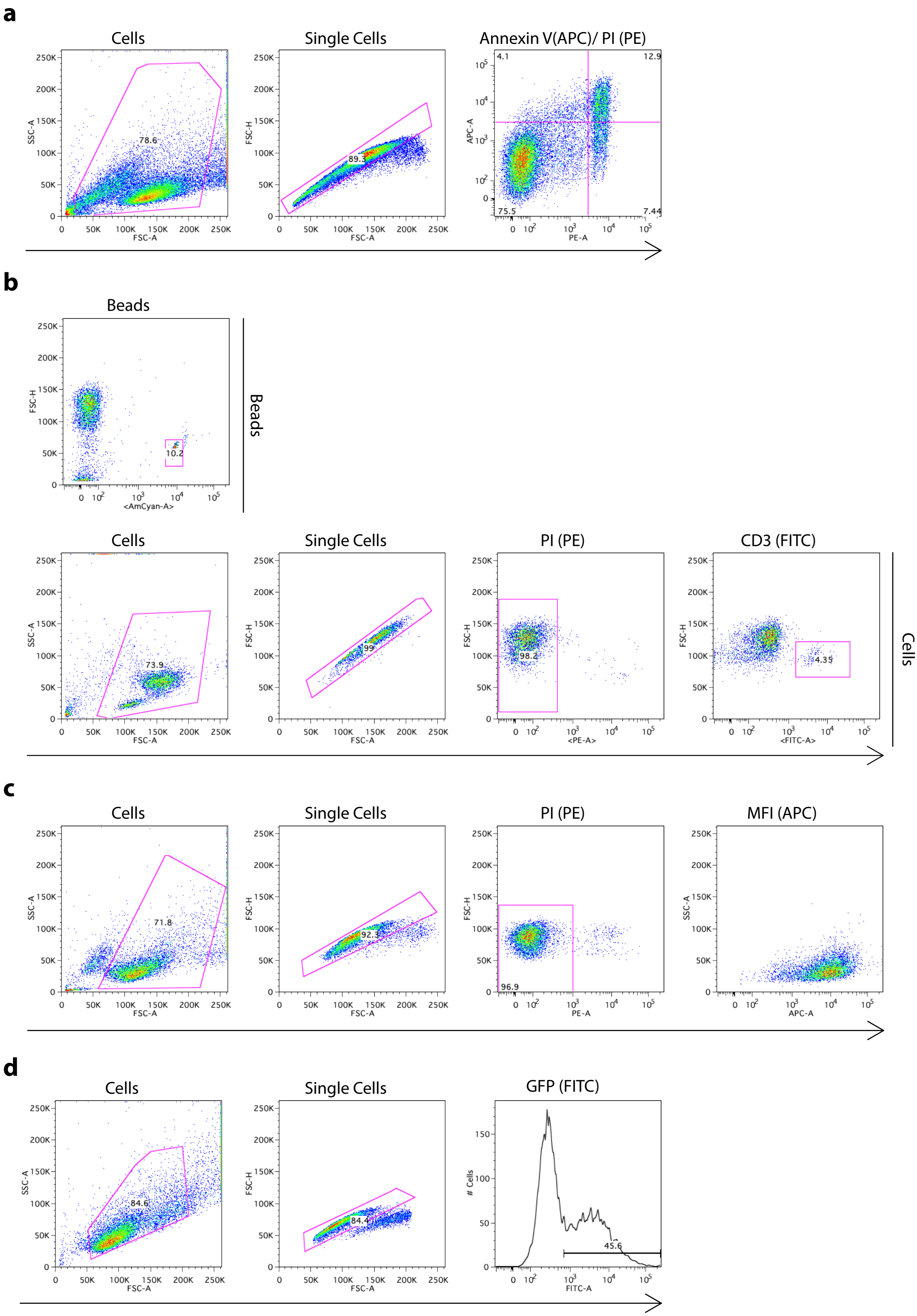
